# QTL Mapping in Outbred Tetraploid (and Diploid) Diallel Populations

**DOI:** 10.1101/2020.12.18.423479

**Authors:** Rodrigo R. Amadeu, Patricio R. Munoz, Chaozhi Zheng, Jeffrey B. Endelman

**Author notes:** Corresponding author: Jeffrey B. Endelman, University of Wisconsin–Madison, 1575 Linden Dr., Madison, WI 53706.

## Abstract

Over the last decade, multiparental populations have become a mainstay of genetics research in diploid species. Our goal was to extend this paradigm to autotetraploids by developing software for quantitative trait locus (QTL) mapping in connected F1 populations derived from a set of shared parents. For QTL discovery, phenotypes are regressed on the dosage of parental haplotypes to estimate additive effects. Statistical properties of the model were explored by simulating half-diallel diploid and tetraploid populations with different population sizes and numbers of parents. Across scenarios, the number of progeny per parental haplotype (pph) largely determined the statistical power for QTL detection and accuracy of the estimated haplotype effects. Multi-allelic QTL with heritability 0.2 were detected with 90% probability at 25 pph and genome-wide significance level 0.05, and the additive haplotype effects were estimated with over 90% accuracy. Following QTL discovery, the software enables a comparison of models with multiple QTL and non-additive effects. To illustrate, we analyzed potato tuber shape in a half-diallel population with 3 tetraploid parents. A well-known QTL on chromosome 10 was detected, for which the inclusion of digenic dominance lowered the Deviance Information Criterion (DIC) by 17 points compared to the additive model. The final model also contained a minor QTL on chromosome 1, but higher order dominance and epistatic effects were excluded based on the DIC. In terms of practical impacts, the software is already being used to select offspring based on the effect and dosage of particular haplotypes in breeding programs.

## INTRODUCTION

For over three decades, the genetic mapping of quantitative trait loci (QTL) using DNA markers has been essential to basic and applied research. Early studies in plants focused on experimental diploid populations created from two parents (Lander and Botstein 1989; Knapp *et al*. 1990). For inbred parents, genetic marker alleles are easily coded to uniquely identify the two parental haplotypes at each locus (Young and Tanksley 1989). For two outbred parents of ploidy *ϕ*, there are 2*ϕ* parental haplotypes (or homologs), but in the case of biallelic SNPs there are only two marker alleles. This difficulty was initially circumvented by using single-dose markers to create separate maternal and paternal maps in diploid species (Grattapaglia and Sederoff 1994) and separate maps for each homologous chromosome in polyploids (Wu *et al*. 1992). Advances in estimation theory (Ritter and Salamini 1996; Wu *et al*. 2002) and software (Van Ooijen and Voorrips 2001; Margarido *et al*. 2007) eventually made it possible to reconstruct F1 diploid offspring in terms of their parental haplotypes, and QTL mapping soon followed (Van Ooijen 2004). Inference of parental haplotypes in F1 polyploid offspring based on bi-allelic SNPs is even more challenging, but several software packages are now available for linkage and QTL analysis (Hackett *et al*. 2017; Bourke *et al*. 2018; Mollinari and Garcia 2019; da Silva Pereira *et al*. 2020).

Not long after QTL mapping in biparental populations became a reality, several groups explored the use of multiple, connected biparental populations via theory and simulation (Rebai and Goffinet 1993; Muranty 1996; Liu and Zeng 2000). We use the term diallel to represent a general class of mating designs in which groups of offspring are derived from no more than two founders (Fig. 1). The transition from theory to practice for diallel mapping began in maize with 25 diverse inbred lines mated to a common parent (Yu *et al*. 2008; Buckler *et al*. 2009), and similar efforts have been published for other crops (Nice *et al*. 2017; Song *et al*. 2017). Examples of QTL mapping in connected F1 populations derived from outbred diploids have also been published (Rosyara *et al*. 2013; Bink *et al*. 2014). Compared with MAGIC (multiparent advanced generation intercross) designs, in which offspring contain haplotypes from more than two founders (Zhang *et al*. 2014; Huang *et al*. 2015), diallel designs have the advantage of mimicking how plant breeding programs typically work. This is particularly important for breeding programs with limited resources to create specialized populations for discovery research.

**Figure 1.**
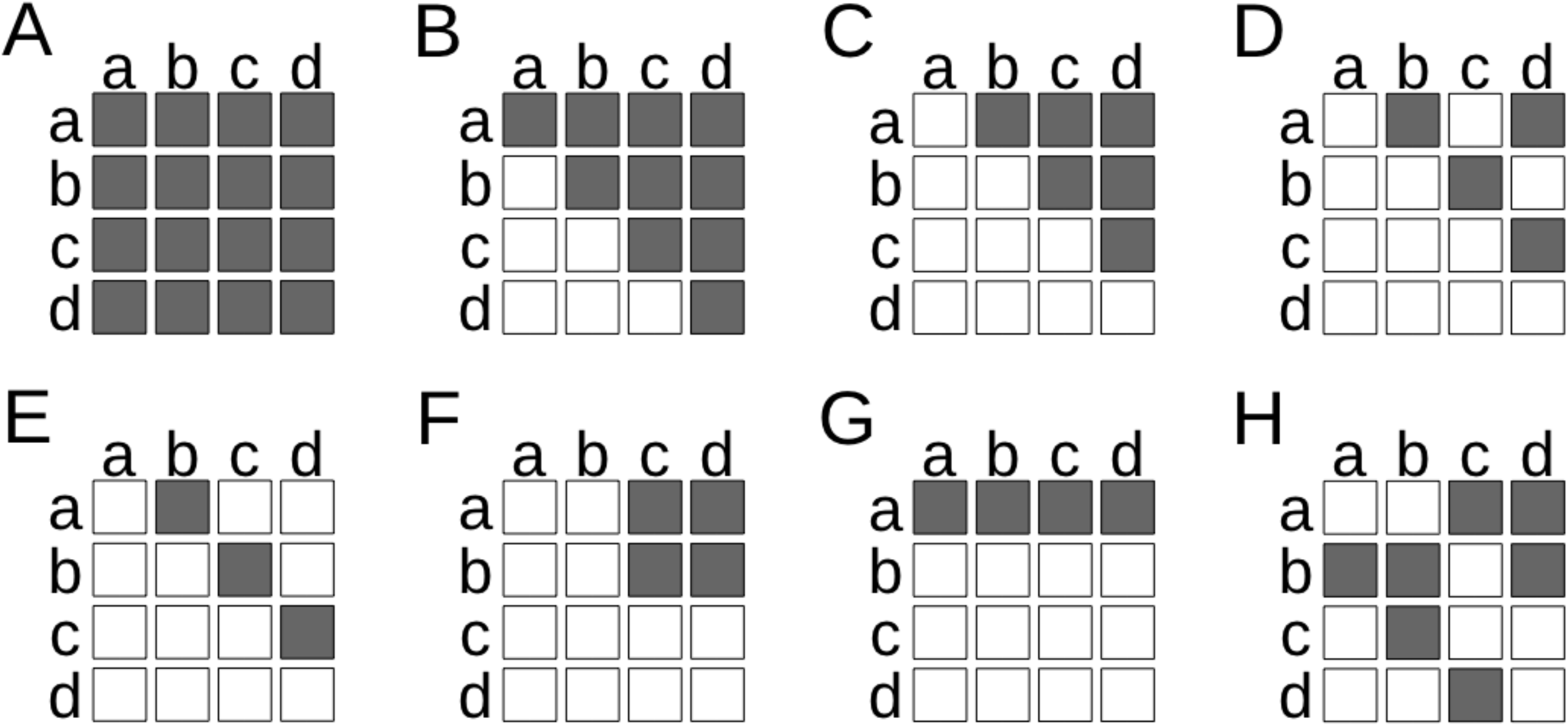
Examples of diallel populations with four parents (labeled a,b,c,d); each square represents a family of full-sibs. (A) full diallel; (B) half diallel with selfing; (C) half diallel without selfing; (D) circular; (E) linear; (F) factorial; (G) testcross or nested design; (H) arbitrary partial diallel. Figure adapted from Verhoeven *et al*. (2006).

The objective of this research was to develop and apply software for QTL mapping in autotetraploid diallel populations (i.e., connected F1 populations), for which no other software is available. Our software, named diaQTL and available as a package for the R Computing Environment (R Core Team 2020), estimates additive effects by regression of phenotypes on the dosage of parental haplotypes. For a diallel with *p* tetraploid parents, there are 4*p* parental haplotypes (some of which may contain identical alleles at a given QTL). diaQTL computes parental haplotype dosage based on output from the companion software PolyOrigin, which estimates parental genotype probabilities using hidden Markov models to perform multi-point linkage analysis (Zheng *et al*. 2021). diaQTL can also analyze connected F1 populations from outbred diploid parents, using parental genotype probabilities from the software RABBIT (Zheng *et al*. 2015).

Our software is also unique for its ability to model dominance effects in tetraploid linkage mapping populations. The dominance deviation for one locus is the residual genetic effect with the additive model. In polyploids, a hierarchy of dominance effects can be defined by regression of the dominance deviation on combinations of parental haplotypes (Kempthorne 1957). Digenic dominance effects are the regression coefficients for a pair of haplotypes, or diplotype; trigenic dominance effects are the regression coefficients for triplotypes; etc.

After exploring the statistical properties of QTL mapping in outbred diallel populations via simulation, we illustrate the use of diaQTL to analyze tuber shape in a tetraploid potato diallel with 3 parents.

## MATERIALS AND METHODS

### QTL Model

A mixed model is used to relate phenotypes to the effects of parental haplotypes and potential covariates. For the *t*^th^ measurement (e.g., plot) of clone *i*, the following equation specifies the additive model for *m* QTL in a diallel with *p* parents of ploidy *ϕ* (bold font designates a vector):

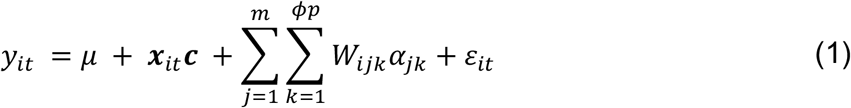

The response variable *y_it_* denotes the phenotype; *μ* is the intercept; ***c*** is an optional column-vector of fixed effects with corresponding row-vector of covariates ***x**_it_*; the additive QTL effects *α_jk_* are random regression coefficients corresponding to the dosage of parental haplotype *k* at locus *j*, denoted *W_ijk_*; and *ε_it_* is the residual.

The dosage of parental haplotype *k* at locus *j*, denoted *W_ijk_*, is calculated from an input file containing the parental genotype probabilities for every offspring. For haplotypes originating in founders that are not the parents of clone *i, W_ijk_* is trivially zero. For the other haplotypes, *W_ijk_* is an expectation over the 100 unique parental genotype states in a biparental tetraploid F1 or 35 unique states in a uniparental (selfed) tetraploid S1 (Zheng *et al*. 2021). If *w_k_* (*abcd*) denotes the dosage of parental haplotype *k* in parental genotype *abcd* (each letter represents a parental haplotype, not necessarily unique), and *P_ij_*(*abcd*) is the genotype probability for clone *i* at locus *j*, then

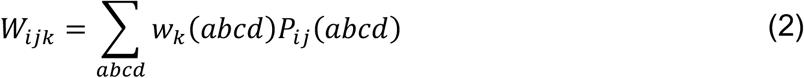

For diploids, the sum in Eq. 2 is over the 4 parental genotype states for a biparental F1 or 3 states for a uniparental S1.

The additive model can be extended to include dominance effects. Denoting the digenic dominance effect for parental haplotypes *k* and *k*′ at locus *j* as *β_jkk′_*, which is a random regression coefficient corresponding to diplotype dosage *W_ijkk′_*, the additive + digenic model is

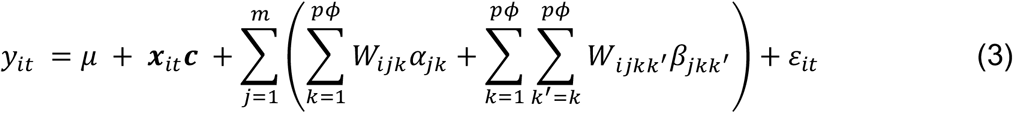

Because the order of the parental haplotypes in diplotype *kk*′ does not matter, the sum over *k*′ in Eq. 3 is restricted to values *k*′ ≥ *k*. For partial diallel designs, not all diplotypes are present, and the sum over *k*′ must be restricted accordingly. Furthermore, digenic dominance may only be included for a subset of the *m* loci based on model selection procedures (see below). The dosage *W_ijkk′_* is computed analogously to Eq. 2, based on the expectation over parental genotypes. For tetraploids, the formula is

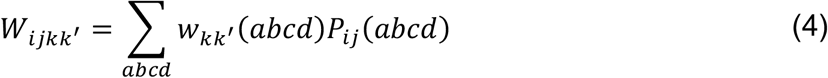

The symbol *w_kk′_*(*abcd*) denotes the dosage of diplotype *kk′* in genotype *abcd*, which equals the number of times *kk′* occurs in the 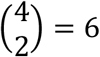 possible combinations of length 2 (sampling from *abcd* without replacement). The definition of higher order dominance effects is analogous to the digenic case and therefore not explicitly shown.

Additive x additive epistatic effects can also be modeled. Denoting the epistatic effect for parental haplotypes *k* and *k′* at loci *j* and *j′*, respectively, as (*αα*)_*jkj′ k′*_, the extension of Eq. 1 is

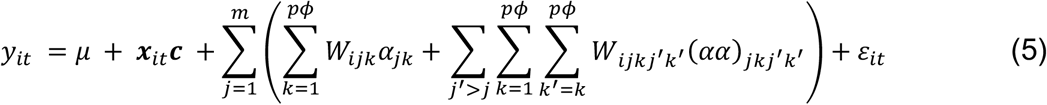

Typically, epistasis is not modeled for all pairs of loci (model selection procedures are used), so limits for the sum over *j*′ are not explicit. The dosage *W_ijkj′ k′_* of the two-locus diplotype is based on the expectation over the parental genotypes at both loci (designated *abcd* and *a′ b′ c′ d′*, respectively):

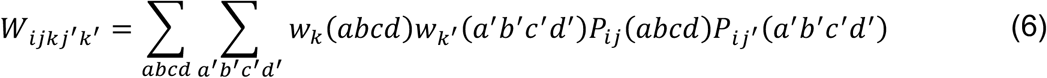

The meaning of the symbols in Eq. 6 is the same as in Eq. 2.

### Polygenic Effects

Polygenic effects can be added to the QTL model for improved understanding of trait genetic architecture and genomic prediction (latter is beyond the scope of this manuscript). Each hierarchy of QTL effects discussed above has an analogous polygenic effect with covariance based on identity-by-descent (IBD) relationships, which can be computed using diaQTL function *IBDmat*.

The additive relationship *A_ii′_* equals ploidy times kinship, which is the probability that a randomly chosen homolog in clone *i* is IBD at a particular locus to a randomly chosen homolog in clone *i′* (Gallais 2003). For locus *j*, the probability that both randomly chosen homologs are parental haplotype *k* is 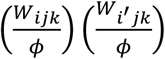, and thus the total probability of IBD (for any parental haplotype) is 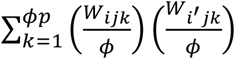. Averaged over *m* loci, the additive relationship becomes

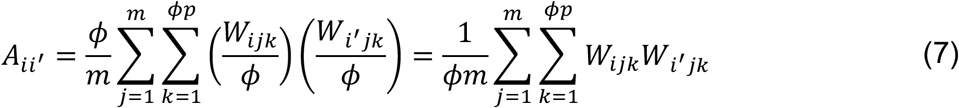

Eq. 7 is computed for each chromosome, and the value used for the polygenic effect is based on averaging all chromosomes that do not contain a QTL; in other words, by extending the leave-one-chromosome-out concept of Yang *et al*. (2014).

IBD relationships for dominance and epistatic effects can be defined analogously to additive relationship (Gallais 2003). The digenic dominance relationship *D_ii′_* equals the binomial coefficient 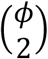 times the probability that a randomly chosen pair of homologs in clone *i* are IBD at a particular locus to a randomly chosen pair in clone *i′*. For locus *j*, the probability that both randomly chosen homologs are parental diplotype *kk′* is 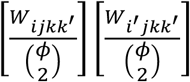, which leads to the following expression:

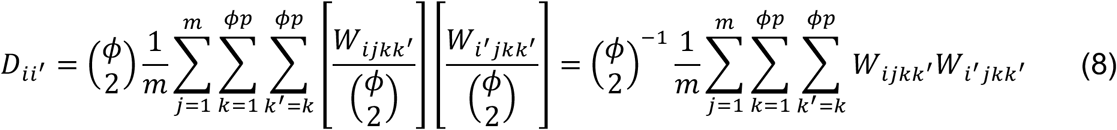

Trigenic and quadrigenic dominance relationships are defined analogously. The additive x additive epistasis relationship *E_ii′_* is also based on IBD probabilities for diplotypes, but in this case the two haplotypes are at different loci (*j* and *j′*), so the probability that both diplotypes are *jkj′ k′* is 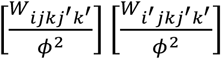 (using the same notation as Eq. 5). The epistatic relationship is based on the average over *ν* pairs of loci from different chromosomes:

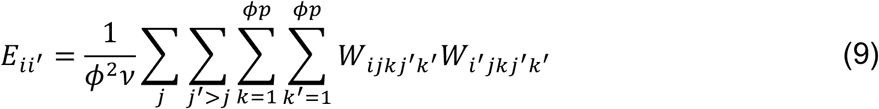

### Bayesian Implementation

R package BGLR (Pérez and de los Campos 2014) was used to implement the regression model due to its flexibility for modeling random effects and its ability to handle binary phenotypes with a generalized linear model (in which case the left-hand side of Eq. 1 is the linear predictor, not the phenotype). BGLR uses a Bayesian framework and Markov Chain Monte Carlo (MCMC) to generate samples from the posterior density. The additive haplotype effects for each QTL (Eq. 1) are independent and identically distributed (i.i.d.) with a “BayesC” prior (Habier *et al*. 2011), which is a two-component mixture of a normal and Dirac distribution (i.e., point mass). This prior was chosen because it accommodates a greater range of complexity for the QTL allelic series than a single normal. The mixing probability and variance of the normal distribution are random (hyper)parameters with their own prior densities (beta and inverse chi-squared, respectively), chosen according to the default rules implemented in the BGLR software. The digenic dominance effects for each locus are i.i.d. with a BayesC prior (separate from the prior for the additive effects), and the same holds for higher order dominance or epistatic effects. The prior density for the additive polygenic effect is multivariate normal with covariance matrix **A** × *V_poly_* (“RKHS” model in BGLR), and the variance component *V_poly_* has an inverse chi-squared prior. The residuals are i.i.d. normal with variance *V_ε_*, which has an inverse chi-squared prior.

If the QTL effects were modeled as fixed, Eq. 1 would be overparameterized, and constraints would be needed to ensure estimability (Hackett *et al*. 2014). Although constraints are not theoretically needed for random effects, in practice we observed large variation in the sum of the additive effects during the Markov chain, even as the differences between the additive effects remained fairly stable. Because ultimately it is these differences that are biologically meaningful, and to reduce the Bayesian credible interval (described below) for the genetic effects, a constant was added to all additive effects (separately for each locus) at each iteration of the Markov chain to impose the constraint of zero sum, and the model intercept was adjusted accordingly. Each group of non-additive effects was similarly constrained to have zero sum.

QTL genetic variances were calculated for each iteration of the Markov chain from the corresponding genetic effects. If 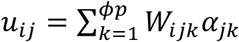 denotes the additive value for clone *i* due to locus *j*, the additive variance for that locus is 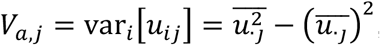, and non-additive variances were computed analogously. The total genetic variance for locus *j*, denoted *V_Q,j_*, is the sum of additive and non-additive variances. The proportion of variance due to the additive effects for locus *j* is

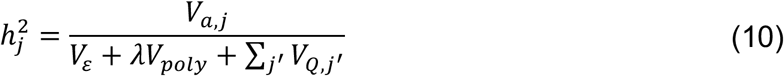

The parameter *λ* in Eq. 10 equals the mean diagonal of the additive relationship matrix, to account for inbreeding in selfed populations (Endelman and Jannink 2012). The proportion of variance due to non-additive or polygenic effects is calculated analogously to Eq. 10.

Two kinds of Bayesian credible interval (CI) are computed in diaQTL. The CI for model parameters is based on the quantiles of the Markov chain (after discarding the burn-in iterations). The CI for QTL location uses a profile likelihood based on the methodology in the R/qtl package (Broman and Sen 2009). Let *f_k_* = *b*(*e^LL_k_^*)*d_k_* be the probability of QTL location being in marker-bin *k*, where *LL_k_* is the posterior mean of the log-likelihood, *d_k_* is the bin-width in cM, and the normalization constant *b* is chosen such that 1 = Σ_*k*_ *f_k_*. The cumulative distribution at bin *k* is 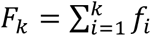. The lower bin for the CI with probability 1 – *α* is one less than the largest *k* satisfying *F_k_* ≤ *α*/2, and the upper bin is one more than the smallest *k* satisfying *F_k_* ≥ 1 – *α*/2.

The burn-in and total number of iterations for MCMC were determined using the Raftery and Lewis (1992) diagnostic implemented in R package coda (Plummer *et al*. 2006). This diagnostic is based on estimating quantile *q* of a parameter within the interval (*q-r, q+r*) with probability *s* (= 0.95 in our analysis). diaQTL function *set_params* returns the value of the diagnostic for each of the genetic and residual variances, and the number of iterations was chosen based on the largest value. For QTL discovery with a one-dimensional scan, which relies on the posterior mean (see below), we used *q* = 0.5, *r* = 0.1. For estimating model parameters and the 90% CI at discovered QTL, we used *q* = 0.05, *r* = 0.025.

### Model Selection

In accordance with the statistical principle of parsimony, we used the Deviance Information Criterion (DIC) to guide the selection of genetic models (Lenarcic *et al*. 2012; Pérez and de los Campos 2014). DIC uses the posterior mean deviance (−2 × log likelihood) to measure model fit and then adds a penalty for model complexity, which equals the difference between the posterior mean deviance and the deviance evaluated at the posterior mean (Spiegelhalter *et al*. 2002). The diaQTL package utilizes DIC for both QTL discovery (with function *scan1*) and post-discovery exploration of multiple QTL and non-additive models (with function *fitQTL*).

For QTL discovery, we used the conventional Neyman-Pearson hypothesis testing framework, with the additive QTL model (Eq. 1) as the alternative hypothesis. The null hypothesis follows Eq. 1, but instead of additive QTL effects, there is one effect for each parent to model general combining ability (GCA). The GCA effects have i.i.d. normal priors, with an inverse chi-squared prior for the variance (“BRR” model in BGLR). Although GCA effects are not explicit in Eq. 1, they are implicitly present because the average of the additive haplotype effects for each parent equals its GCA.

The test statistic (i.e., “score”) for each marker bin during the genome scan is −ΔDIC, which equals the DIC of the null hypothesis minus the DIC of the alternative hypothesis. A typical convention for accepting a more complex model is that the DIC should decrease by at least 5, preferably 10 (Lunn *et al*. 2012), but this is not adequate to control the genome-wide Type I error rate during QTL discovery. To determine a proper threshold, one can use diaQTL function *scan1_permute* to run a stratified permutation test (Churchill and Doerge 1994), randomly permuting the phenotypes within each F1 population. The largest −ΔDIC for each permutation is recorded, and the 1 – *α* quantile of this distribution is used for QTL discovery with significance level *α*.

Alternatively, due to the generic nature of linkage mapping populations for a given mating design, number of parents, ploidy, and genome size, simulation can be used to determine the appropriate threshold. Half-diallel mating designs (without selfing) were simulated using the software PedigreeSim V2.0 (Voorrips and Maliepaard 2012) and auxiliary R package PedigreeSimR (https://www.github.com/rramadeu/PedigreeSimR). The simulated genomes contained 1 to 12 linkage groups, each with a length of 100 cM, loci evenly spaced at 0.1 cM, and recombination based on Haldane’s map function. For tetraploids, only bivalents were allowed with no preferential pairing of homologous chromosomes. The number of parents was varied from 2 to 10. A genome-wide scan with *scan1* was run for 1000 simulations, using standard normal deviates as phenotypes, and the largest −ΔDIC for each simulation was recorded. For each ploidy (2x and 4x) and significance level (*α* = 0.01,0.05,0.1,0.2), a bivariate monotone regression spline was fit for genome size and number of parents using R/scam (Pya and Wood 2015). Predicted values from the spline are returned by diaQTL function *DIC_thresh*.

### Power Simulation

A factorial numerical experiment with 3 × 3 × 3 × 2 = 54 treatment scenarios, and 1000 simulations per scenario, was used to estimate statistical power and accuracy as a function of the total population size (*N* = 200, 400, 800), number of parents (3, 4, 6) with a half-diallel mating design, heritability (*h*^2^ = 0.1, 0.2, 0.4), and ploidy (2, 4), assuming a genome size of 12 Morgans (simulated as described above). The −ΔDIC threshold was chosen to maintain significance level *α* = 0.05. Simulations were performed using a high-performance computing facility at the University of Florida (HiPerGator 2.0), allocating 2GB of RAM for each thread.

For each simulation, a single additive QTL was randomly assigned to one locus, and all other loci were used as markers for mapping. Unreplicated phenotypes were simulated by 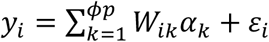 (same symbols as Eq. 1), using i.i.d. standard normal deviates for the additive haplotype effects and i.i.d. normal residuals with variance chosen to achieve the target heritability. If the realized heritability, computed from the simulated QTL effects and residuals (Eq. 10), deviated from the prescribed value by more than 0.01 (due to finite sample size), the simulation was discarded before QTL mapping. Perfect knowledge about the haplotype dosage *W_ik_* was assumed, which is asymptotically achievable with PolyOrigin as marker density increases (Zheng *et al*. 2021). Genotyping error would reduce power and accuracy (Zhang *et al*. 2014; Bourke *et al*. 2019).

Statistical power was the proportion of simulations in which both flanking markers of the simulated QTL were declared significant. The accuracy of the inferred QTL position was the distance between the simulated position and most significant marker. Accuracy for the posterior mean estimates of the additive effects was evaluated based on the Pearson correlation with simulated values. Monotone splines for power and accuracy were fit using R/scam (Pya and Wood 2015).

To ensure the results were robust to polygenic effects, we simulated traits with an additive QTL and polygenic effect for a half-diallel design with 4 parents and total population size *N* = 200. The multivariate normal deviate was simulated using *mvrnorm* from R/MASS (Venables and Ripley 2002), with covariance equal to the additive relationship matrix (Eq. 7) times a polygenic variance component. Increasing the proportion of variance due to the polygenic effect up to 0.3, while keeping the QTL h^2^ at 0.2, had negligible effect on statistical power (Fig. S1).

### Potato Data

The potato dataset is a half-diallel population (without selfing) for three parents from the University of Wisconsin (UW) breeding program: Villetta Rose, W6511-1R, and W9914-1R. The population was genotyped using version 3 of the potato SNP array, and allele dosage was called using R package *fitPoly* (Voorrips *et al*. 2011; Zych *et al*. 2019). There were 5334 markers distributed across 12 chromosome groups, and their physical positions were based on the potato DMv4.03 reference genome (Potato Genome Sequencing Consortium 2011; Sharma *et al*. 2013). Parental genotype probabilities and a genetic map were calculated using the software PolyOrigin (Zheng *et al*. 2021), which identified 19 of the 434 progeny as outliers, and phenotype data were unavailable for 2 more. The remaining 413 clones were distributed as follows: Villetta Rose × W6511-1R (154 individuals), Villetta Rose × W9914-1R (113 individuals), and W6511-1R × W9914-1R (146 individuals). The genetic map produced by PolyOrigin spanned 12.1 Morgans and contained 2781 marker-bins.

The half-diallel population was evaluated as part of a larger field trial in 2018 at the UW Hancock Agricultural Research Station. The trial used an augmented design with four incomplete blocks and seven repeated checks per block. Each plot was a single row sown with 15 seeds (tuber pieces) at 30 cm in-row spacing and 90 cm between rows. The trial was harvested mechanically, and tubers were passed through an optical sizer after washing to measure tuber length and width. Tuber shape was calculated as the average length/width (L/W) ratio of all tubers weighing 170–285g. To improve the normality of the residuals, the trait was transformed to log [(*L/W*) – 1]. Initial statistical analysis using ASReml-R v4 (Butler *et al*. 2018) was based on the linear model *y_ij_* = *μ* + *block_j_* + *clone_i_* + *ε_ij_*. Broad-sense heritability on a plot basis was estimated at 0.91 by treating *clone_i_* and *block_j_* as random effects (i.i.d. normal). BLUEs for each clone were estimated by treating eZone# as fixed and then used as the response variable for analysis with diaQTL.

To give an idea about computational requirements, a variable of size 430 MB was created by the diaQTL function *read_data* from the potato input CSV files. Based on the output from *set_params*, 500 iterations were used with *scan1* for QTL discovery, which required 3 minutes using 2 cores of a 3.1 GHz Intel Core i5 processor. Several functions in the diaQTL package (*read_data, scan1, IBDmat*) can utilize multiple cores for parallel execution.

### Software and Data Availability

Version 1.00 of diaQTL and version 0.2 of PedigreeSimR, which were current at the time of manuscript publication, are available as supplemental files through FigShare under the GPL-3 license. For the most up-to-date version of diaQTL, download the package from https://github.com/jendelman/diaQTL. The potato dataset is distributed with the diaQTL package and used in the tutorial vignette.

## RESULTS

### Statistical power and accuracy

Simulated half-diallel populations were generated to study the influence of ploidy, number of parents, and genome size on the threshold needed to control the genome-wide Type I error rate at *α* = 0.05. The test statistic is −ΔDIC, which equals the Deviance Information Criterion for the null hypothesis (GCA but no QTL effects) relative to the alternative hypothesis of an additive QTL. The threshold was higher for tetraploids compared to diploids and increased approximately linearly with the number of parents (Fig. 2). There was also an approximately linear relationship between the threshold and logarithm of genome size (Fig. S2). As expected from previous research in diploids (Lander and Botstein 1989), the threshold was not influenced by population size. QTL discovery can also be conducted using markers as covariates or with digenic dominance, but higher −ΔDIC thresholds are needed to maintain the same significance level (Fig. S3).

**Figure 2.**
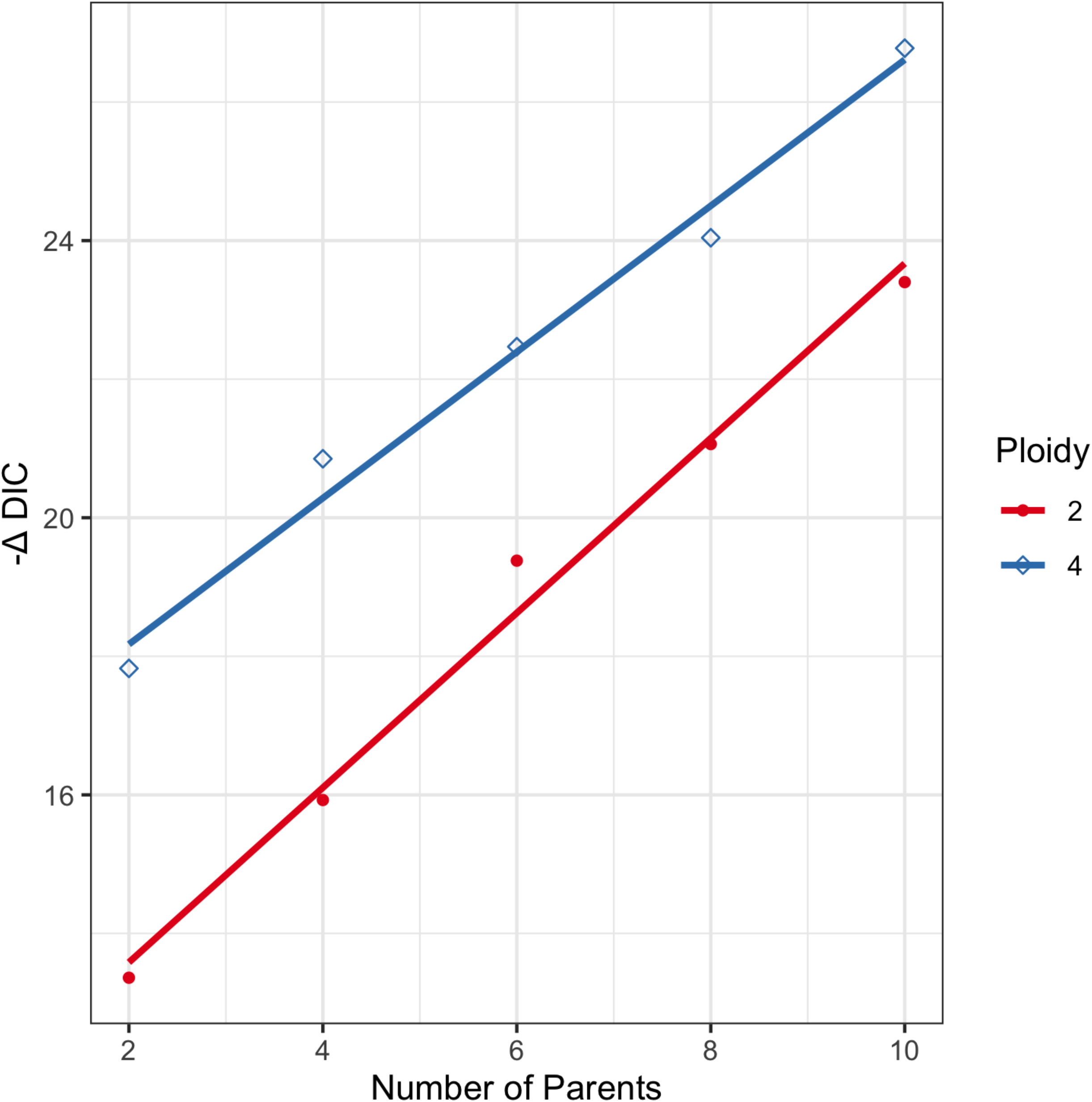
Threshold to control the genome-wide Type 1 error rate at α = 0.05, for a half-diallel design without selfing. The statistic −ΔDIC is the Deviance Information Criterion for the null hypothesis (no QTL) relative to the alternative hypothesis of an additive QTL. Each point is based on 1,000 simulations, and the best-fit linear regression is shown as a solid line.

The influence of population size, number of parents, ploidy, and QTL heritability on statistical power was investigated for a genome of 12 Morgans (the size of the potato genome). Power increased with population size and QTL heritability but decreased with the number of parents and ploidy (Fig. S4). For a given h^2^, the interplay between these factors could be largely summarized by the number of progeny per parental haplotype, abbreviated pph, but the diploid (tetraploid) results were consistently below (above) the regression spline (Fig. 3 and S5). At h^2^ = 0.1 and α = 0.05, power reached 0.5 and 0.9 with 30 and 70 pph, respectively, while only 10 and 25 pph (respectively) were needed for h^2^ = 0.2. At h^2^ = 0.4, the power was 1 for all scenarios (Fig. S4). As power increased, so did the accuracy of the inferred QTL position, as measured by the distance between the most significant marker and QTL (Fig. S6). The positional accuracy was 6 cM when the power was 0.5 and decreased to 3 cM when power was 0.9.

**Figure 3.**
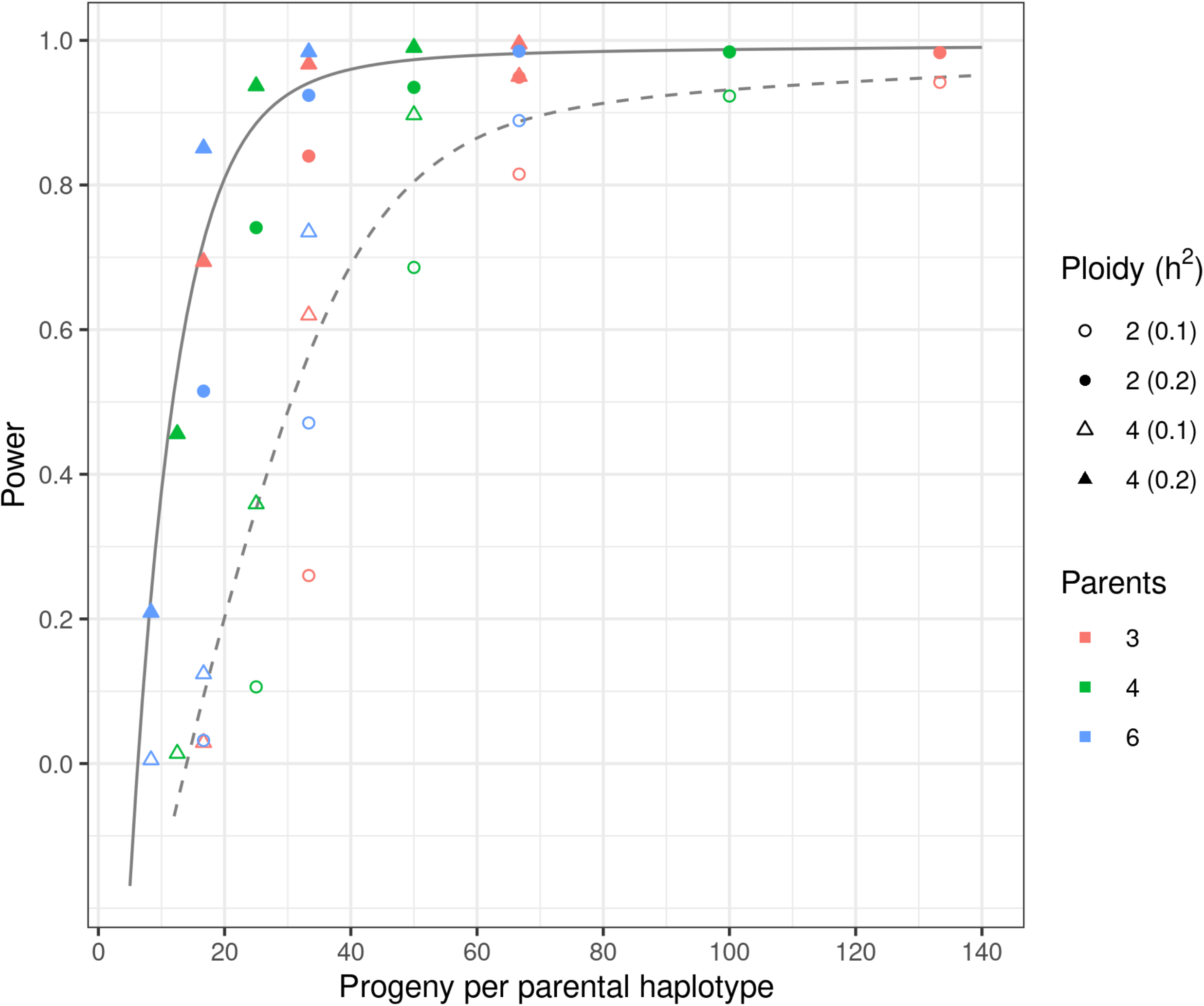
Statistical power to detect a multi-allelic QTL at significance level α = 0.05, as a function of the number of progeny per parental haplotype, number of parents, ploidy, and QTL heritability (*h^2^*). Each point is the average of 1,000 simulations with an additive model. The dashed line is a monotone increasing, concave spline for *h^2^* = 0.1, and the solid line is the spline for *h^2^* = 0.2.

The accuracy of the predicted haplotype effects, as measured by the correlation with simulated values, was also largely explained by the pph metric (Fig. 4). For h^2^ = 0.1 and α = 0.05, 40 pph was sufficient to achieve an accuracy of 0.9, while for h^2^ = 0.2, only 20 pph was needed. Across all scenarios, the heritability of the QTL could be estimated with essentially no error (Fig. S4), even when the accuracy of the haplotype effects was only 0.7.

**Figure 4.**
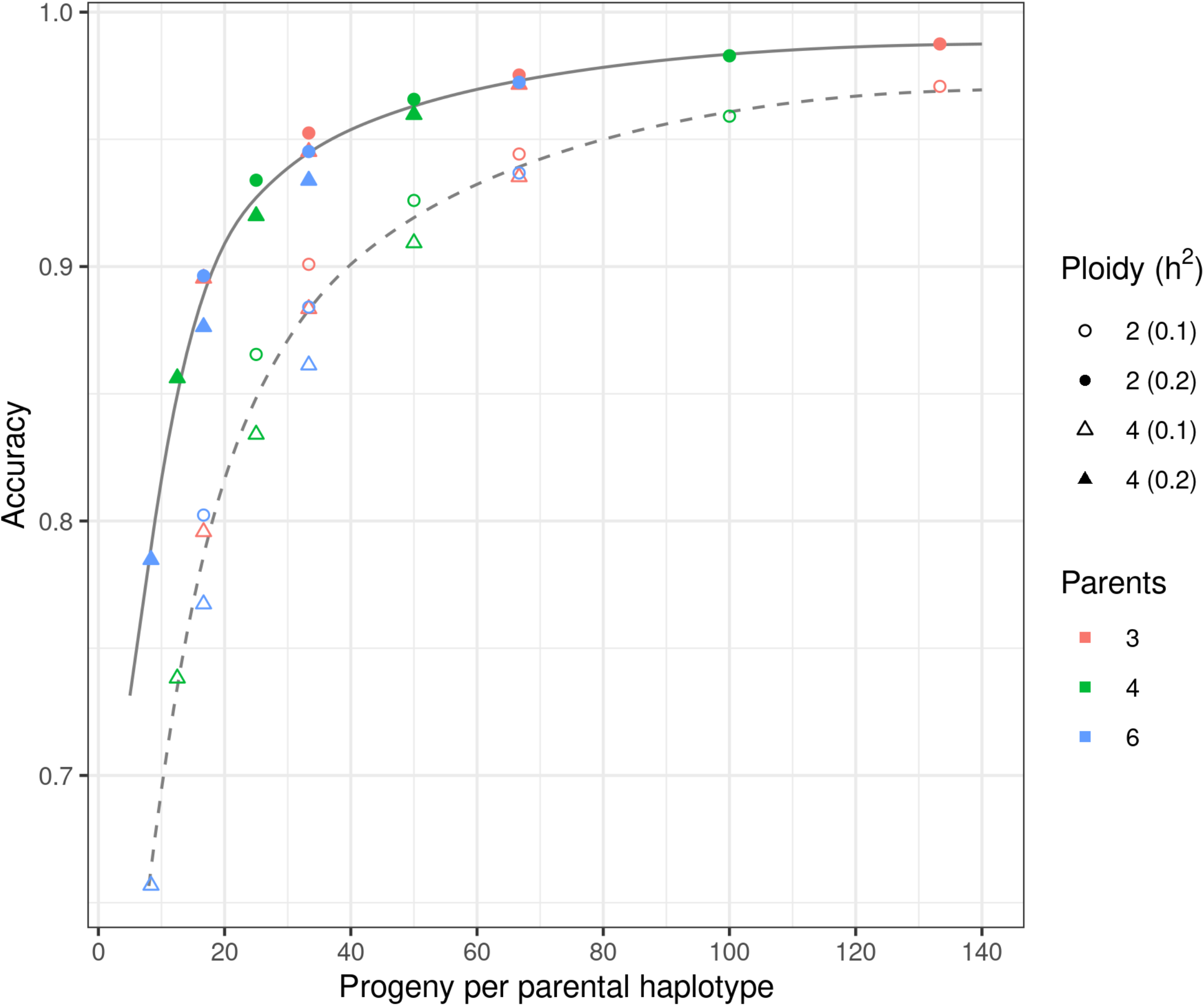
Accuracy of the estimated haplotype effects, as measured by Pearson’s correlation with the simulated values. Each point is the average of 1,000 simulations with an additive model. The dashed line is a monotone increasing, concave spline for *h^2^* = 0.1, and the solid line is the spline for *h^2^* = 0.2.

### Potato diallel

To demonstrate diaQTL with a real dataset, we analyzed potato tuber shape, measured by the length to width (L/W) ratio, in a half diallel population with three parents. The ideal range of values for tuber shape is determined by the end-product (e.g., long tubers for French fries, round tubers for potato chips) and to some extent cultural preferences. The three parents were developed for the red-skin fresh market in the US, for which a round to slightly oblong shape is expected. The distribution of L/W values for the population ranged from 1.0 to 2.0 (Fig. S7), while the commercial checks used in the field experiment had values between 1.1 and 1.2.

A genome scan with the additive model identified a large peak on chromosome 10 at 63 cM (Fig. 5A), designated 10@63. This region coincides with the location of the classical *Ro* (round) QTL in potato (Van Eck *et al*. 1994), and the causal gene has been identified as *StOVP20*, an *OVATE Family Protein* with orthologs that affect fruit shape in tomato, melon, and cucumber (Wu *et al*. 2018). *StOVP20* is not present in the DM reference genome, but based on synteny analysis (Wu *et al*. 2018), it is flanked by the DM gene models (PGSC0003DMG400006678, PGSC0003DMG400006679). This corresponds to the DM genome interval (48,982,521 - 49,022,687), which lies within the 95% CI for the 10@63 QTL (48,203,284 - 50,782,097).

**Figure 5.**
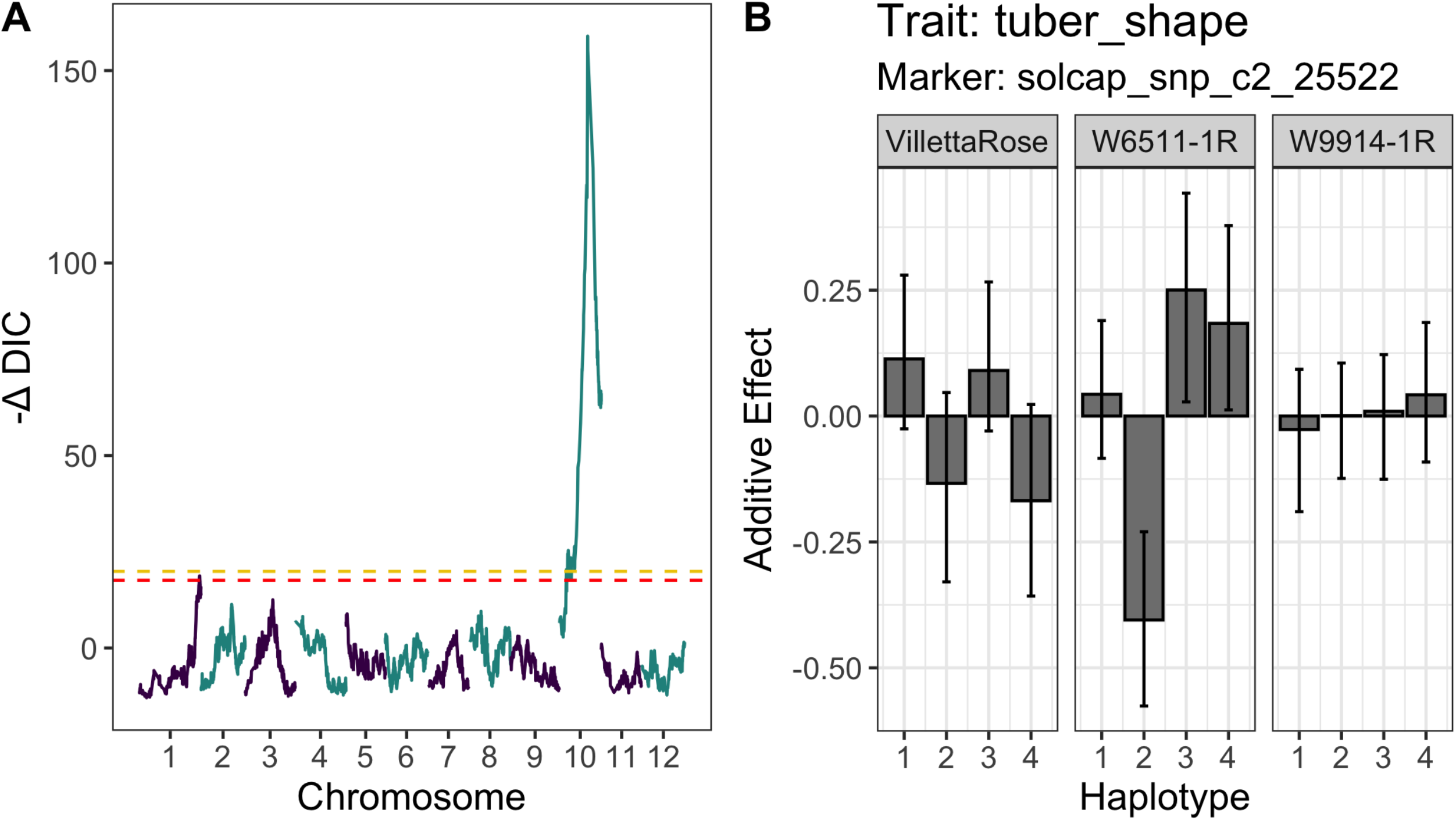
R/diaQTL results for potato tuber shape using a half-diallel population with three parents: VillettaRose, W6511-1R, and W9914-1R. (A) Genome scan with *scan1*. The dashed horizontal lines correspond to α = 0.05 (gold) and α = 0.1 (red). The most significant marker was solcap_snp_c2_25522 on potato chromosome 10. (B) Additive effect estimates for the 12 parental haplotypes using *fitQTL*. Error bars are the 90% CI.

The next highest peak in the genome scan was on chromosome 1 at 133 cM. The −ΔDIC score was at the threshold for α = 0.05 but clearly above the threshold for α = 0.1 (Fig. 5A). The model with both QTL reduced the DIC by 23 points compared to the model with only 10@63, so it was accepted over the single QTL model (Table 1).

**Table 1.** Model comparison for potato tuber shape based on ΔDIC, which is the Deviance Information Criterion relative to a null model with parental GCA effects. QTL are designated as chromosome@cM, with A = additive, D = digenic, T = trigenic, and AA = additive x additive epistasis.

| Model | Effects | ΔDIC |
| --- | --- | --- |
| 1 | 10@63(A) | -160 |
| 2 | 10@63(A) + 1@133(A) | -183 |
| 3 | 10@63(A) + 1@133(A) + 10@63 x 1@133 (AA) | -189 |
| 4 | 10@63(A+D) + 1@133(A) | **-200** |
| 5 | 10@63(A+D) + 1@133(A) + 10@63 x 1@133 (AA) | -196 |
| 6 | 10@63(A+D+T) + 1@133(A) | -197 |
| 7 | 10@63(A+D) + 1@133(A+D) | -196 |

More complex models involving dominance and/or additive x additive epistatic effects were evaluated next (Table 1). Epistasis lowered the DIC by 6 points compared to the two-QTL additive model, whereas digenic dominance at 10@63 lowered the DIC by 17 points. Including both of these non-additive effects raised the DIC, as did higher order dominance at 10@63 and dominance at 1@133 (Table 1). We therefore selected the model with additive and digenic effects at 10@63 and additive effects at 1@133, which accounted for 26%, 8%, and 5% of the total variance, respectively (Table 2). To quantify the importance of smaller undetected QTL, an additive polygenic effect was included, which lowered the DIC by 78 points and accounted for 29% of the total variance (Table 2).

**Table 2.**
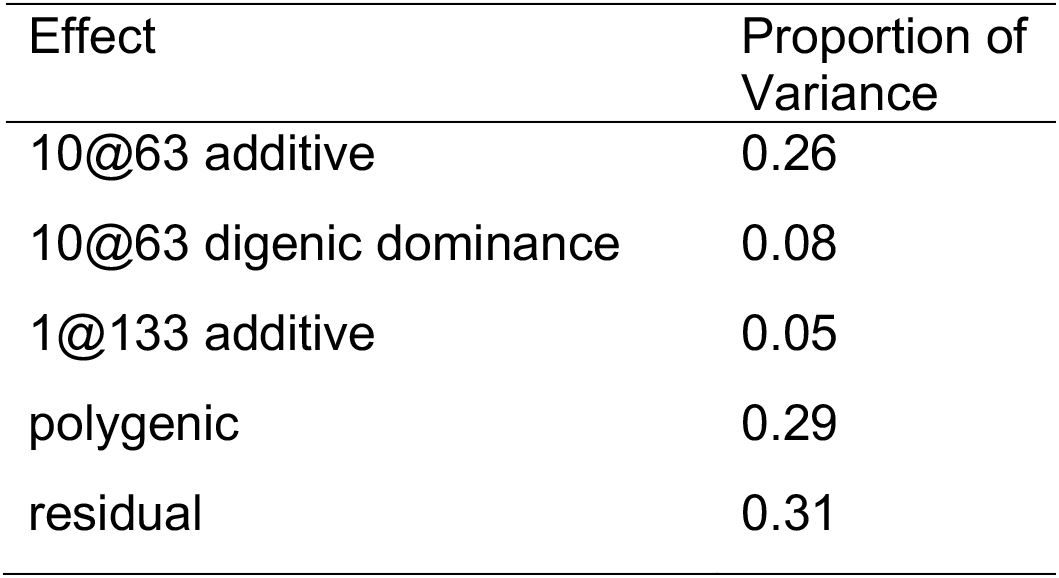
Proportion of variance for potato tuber shape based on Model 4 from Table 1, plus a polygenic effect. QTL are designated as chromosome@cM.

Of the 12 parental haplotypes at 10@63, the largest additive effect (in magnitude) was estimated for haplotype 2 of parent W6511-1R (Fig. 5B), and the negative sign means it reduced the L/W ratio. Haplotypes 2 and 4 of parent Villetta Rose also reduced the L/W ratio. For parent W9914-1R, all four haplotypes had similar additive effects but showed some differentiation with respect to non-additive effects (Fig. S8 and S9).

## DISCUSSION

Although this is the first publication on statistical power and accuracy for QTL mapping in tetraploid diallel populations, some comparisons with previous studies on diploid diallel or biparental tetraploid populations are possible. Among previous simulation studies, Muranty (1996) is unique for explicitly showing the loss of statistical power as the number of outbred diploid parents increased at a fixed total population size. We observed the same trend (Fig. 3), but the exact values for power are not comparable because of differences in the significance level (higher values produce higher power). Muranty (1996) also reported that the specific form of the mating design (e.g., cyclic, factorial, half-diallel) had little impact on power, which was confirmed in our simulation (data not shown). Bourke *et al*. (2019) reported statistical power in a biparental tetraploid population of size 200, which corresponds to 25 pph, was 0.8 for a bi-allelic, additive QTL with h^2^ = 0.1 and high genotypic information content. This is higher than the value of 0.4 for the regression spline in Fig. 3, which may be the result of the simpler genetic architecture (bi-allelic vs. multi-allelic in our simulation).

Compared to existing software, R/diaQTL has unique features for modeling dominance and epistasis with multi-allelic QTL in tetraploids. The inner product of the genetic dosages used for QTL mapping of non-additive effects, when averaged across many loci, generates non-additive relationships (see Methods). Due to Mendelian segregation, these realized relationships exhibit a distribution around the expected value based on pedigree (Fig. S10; Amadeu *et al*. 2020). Non-additive relationship matrices probably have limited utility for QTL mapping, but we envision potential applications with genomic selection. Breeding values include 1/2 of the additive x additive epistasis and 1/3 of the digenic dominance in tetraploids (Gallais 2003), although this result is only exact when genotype frequencies equal the product of haplotype frequencies (Kempthorne 1957), which requires a complete diallel with selfs.

Our statistical model allows each parental haplotype to have a different effect, but we expect some haplotypes will carry the same QTL allele. Although the use of random effects for the regression coefficients helps offset the statistical complexity of the haplotype model, from a genetic point of view, we are interested in inferring identity-by-state relationships. At present, a user of our software is limited to hypothesizing that haplotypes with similar effect carry the same allele. One direction for future research would be to incorporate methods to formally test such hypotheses (Jannink and Wu 2003; Crouse *et al*. 2020).

Large plant breeding programs often maintain distinct research enterprises devoted to genetic discovery vs. variety selection. Genetic discovery projects typically characterize novel populations, such as a panel of distantly related germplasm or larger-than-normal, unselected biparental populations, but this additional effort is not always feasible for small programs. There is also the potential disconnect between the development of a genetic marker in the discovery population and its successful deployment in the breeding populations; such markers often fail because they are not haplotype-specific. With a diallel QTL analysis of the breeding population, this additional step would be unnecessary. The diaQTL function *haplo_plot* allows users to visualize which parental haplotypes are present at a particular locus (Fig. S11), and it is straightforward to select progeny-based on the dosage of particular haplotypes using diaQTL function *haplo_get* (see package tutorial).

The University of Wisconsin potato breeding program provides an example to discuss the feasibility of using breeding populations for discovery research. A typical mating scheme (for each market type) involves 10 females and four males (male fertility is scarce), from which 40 F1 populations with 500 progeny each are created and evaluated as single plants in the field. Using visual selection for plant maturity and tuber type, 800 individuals (= 4%) advance to the first clonal stage, for genotyping and further phenotyping. Under uniform selection, this population size corresponds to 20 pph for the female parents (more for the male parents), which translates into 80% power at h^2^ = 0.2 based on our simulation results.

## CONCLUSION

The R/diaQTL software is a user-friendly tool for QTL mapping and haplotype analysis in connected F1 populations derived from outbred parents. The number of progeny (or gametes, if selfed populations are used) per parental haplotype is a critical design parameter to consider when developing populations. By enabling selection on the effect and dosage of haplotypes without having to develop haplotype-specific markers, diaQTL can improve the efficiency of breeding.

## Acknowledgments

Financial support provided by USDA NIFA Award No. 2019-67013-29166. We thank Maria Caraza-Harter for testing the diaQTL software.

## Author contributions

RRA and JBE developed the software, analyzed the data, and drafted the manuscript. All authors contributed to the methodology and edited the manuscript.

## Supplemental Figures

**Figure S1.**
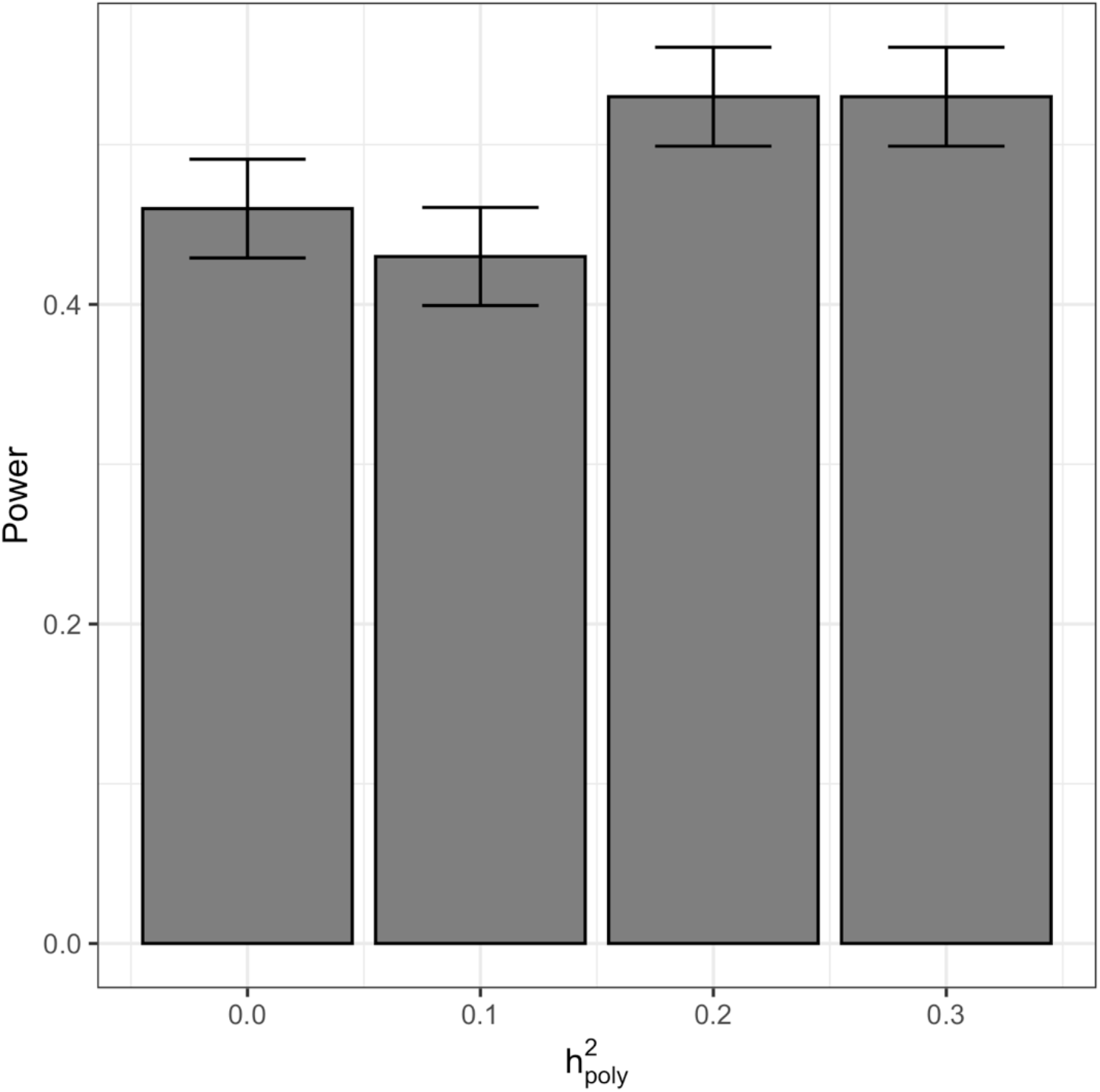
The influence of simulating polygenic effects on statistical power for detecting an additive QTL with h^2^ = 0.2 in a half-diallel with 4 parents and total population size N = 200. The height of each bar is the proportion of 1000 simulations in which the QTL was detected, and the error bars are ±1.96 SE for the standard error (SE) of a binomial proportion.

**Figure S2.**
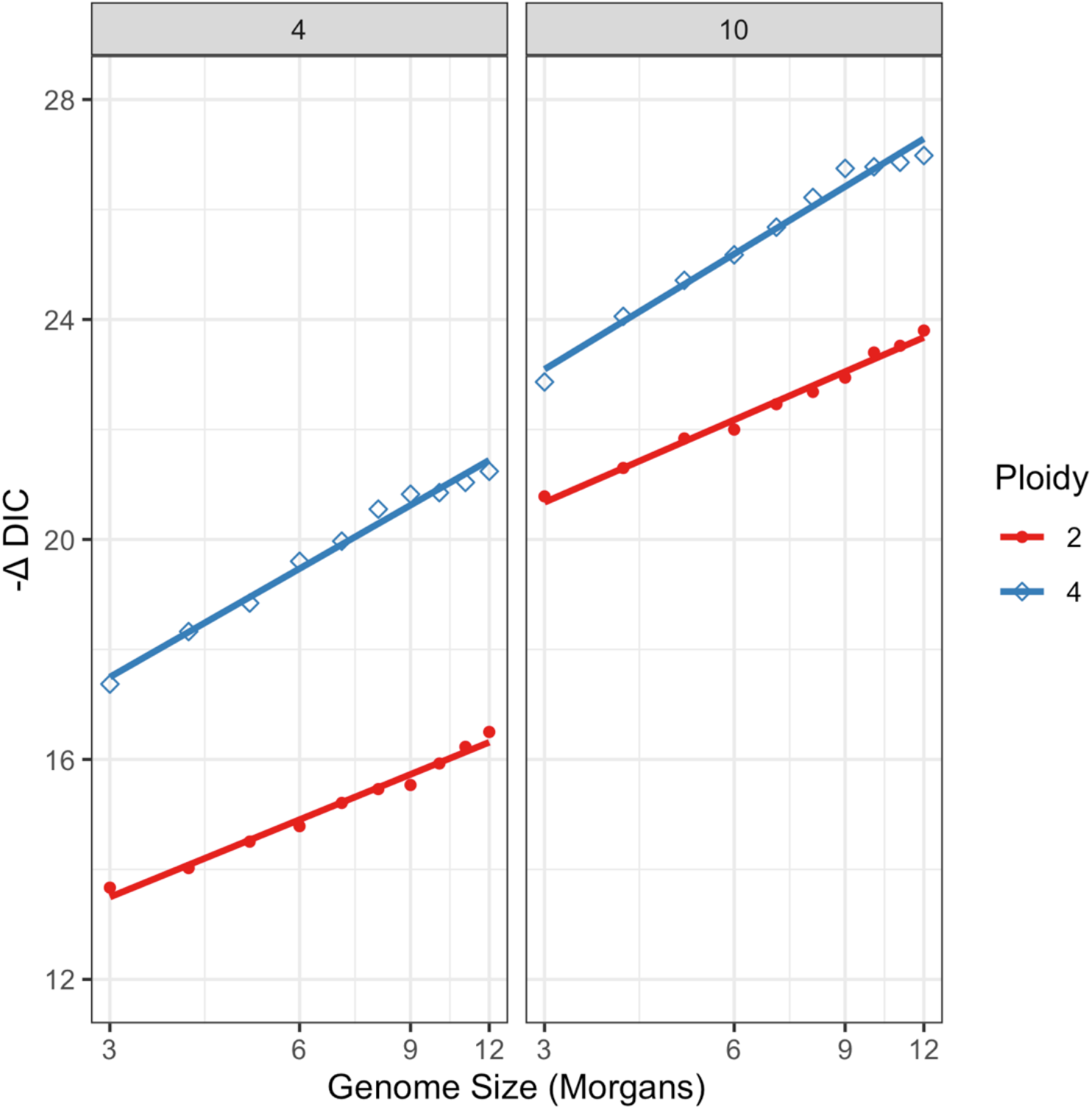
The threshold for QTL discovery, −ΔDIC, increased approximately linearly with the logarithm of genome size. Results are shown for significance level *α* = 0.05, using simulated half-diallel populations, and the number of parents (4 and 10) is shown at the top of each panel. Each point is based on 1000 simulations.

**Figure S3.**
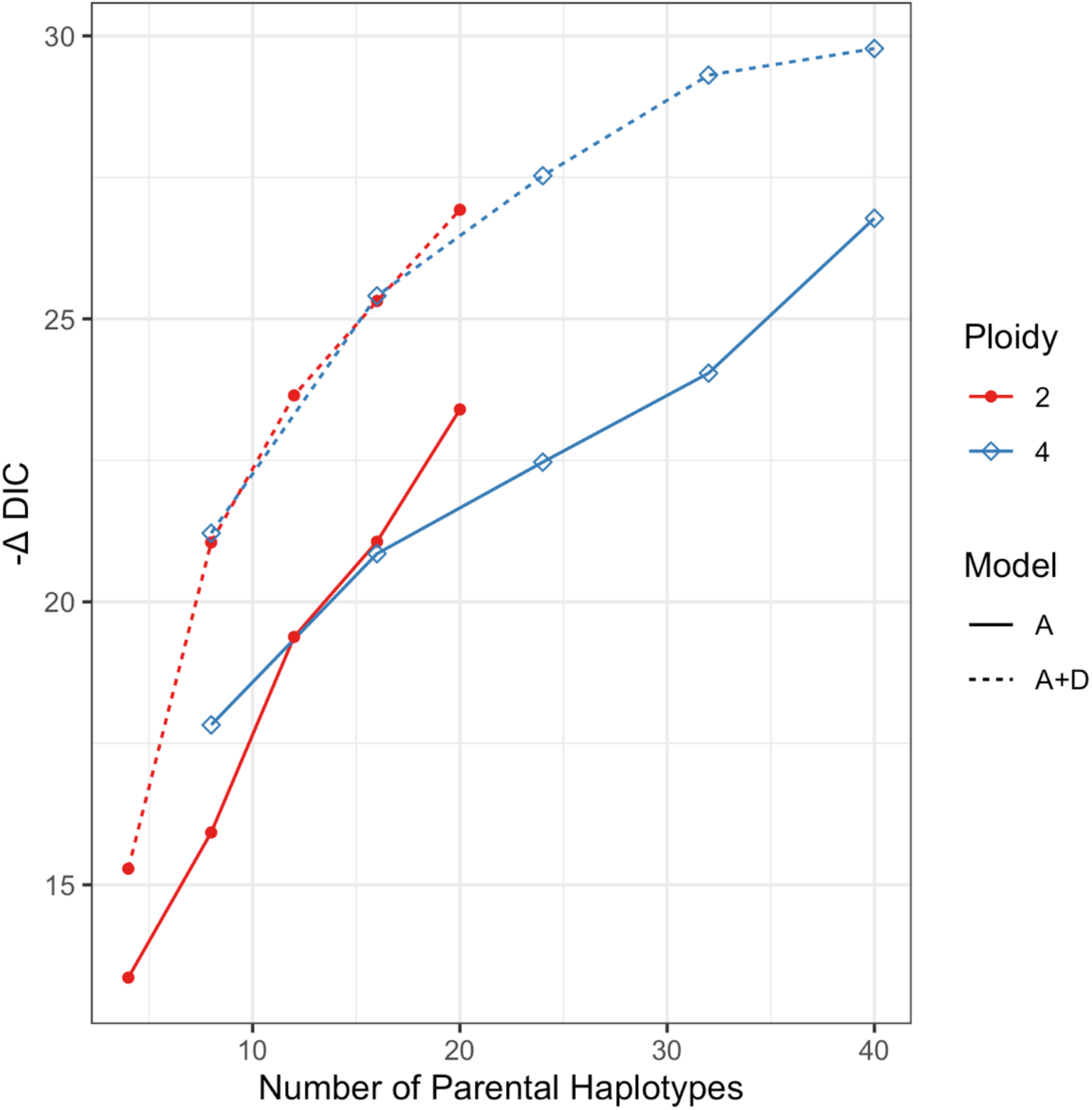
The threshold for QTL discovery, –ΔDIC, was higher for a QTL model with additive and dominance effects (A+D) compared to only additive (A) effects. Results are shown for significance level *α* = 0.05, using 1000 simulated half-diallel populations. The number of parental haplotypes equals ploidy (2 or 4) times the number parents (2 to 10).

**Figure S4.**
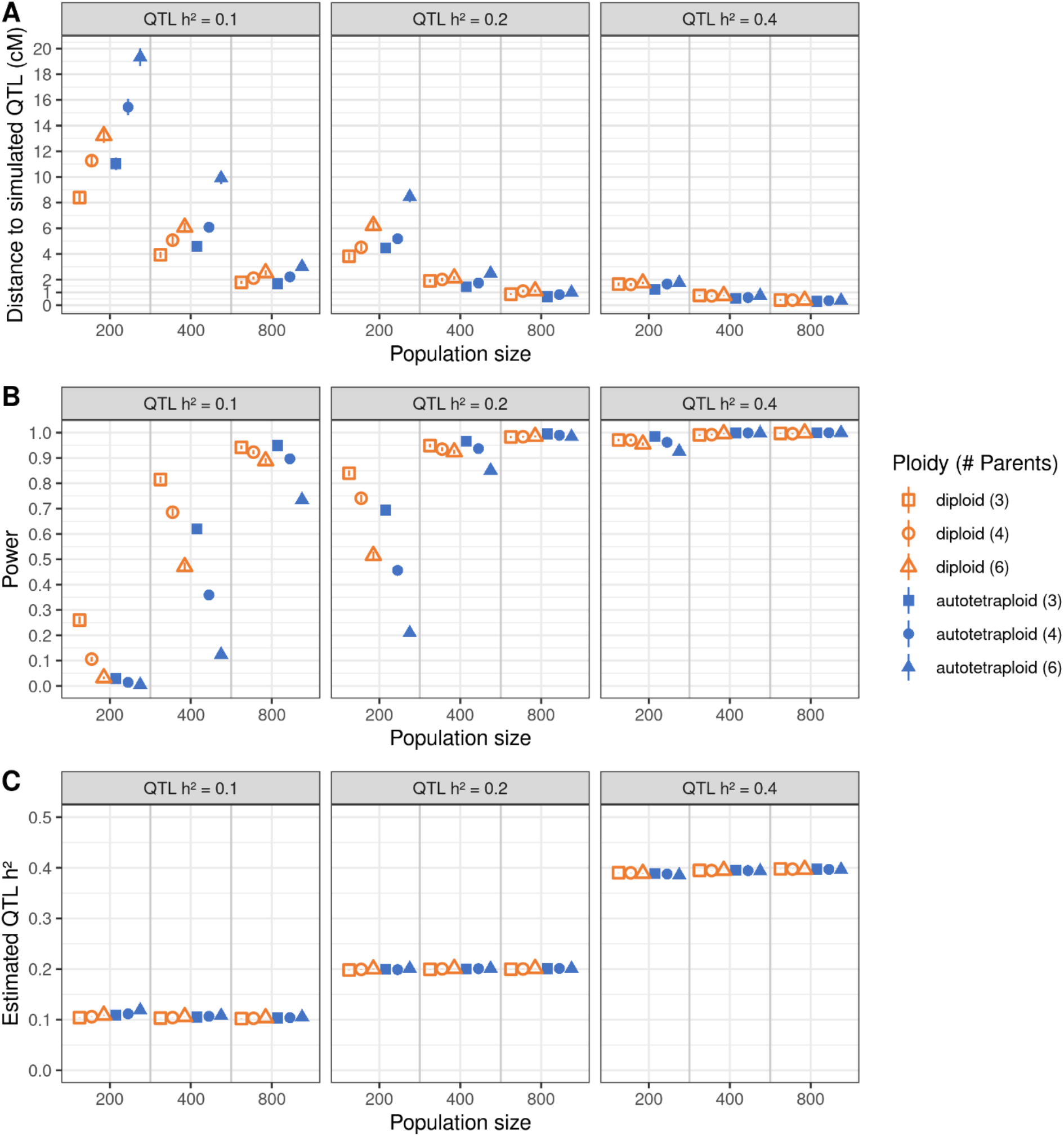
QTL mapping results from simulated half-diallel populations with varying population size, QTL heritability, number of parents, and ploidy. The genome-wide Type I error rate was controlled at *α* = 0.05. A) Average distance in cM between the peak LOD score position and simulated QTL position. B) Average power to detect the QTL. C) Average estimated QTL heritability based on the diaQTL linear model. Error bars are standard errors based on 1,000 simulations.

**Figure S5.**
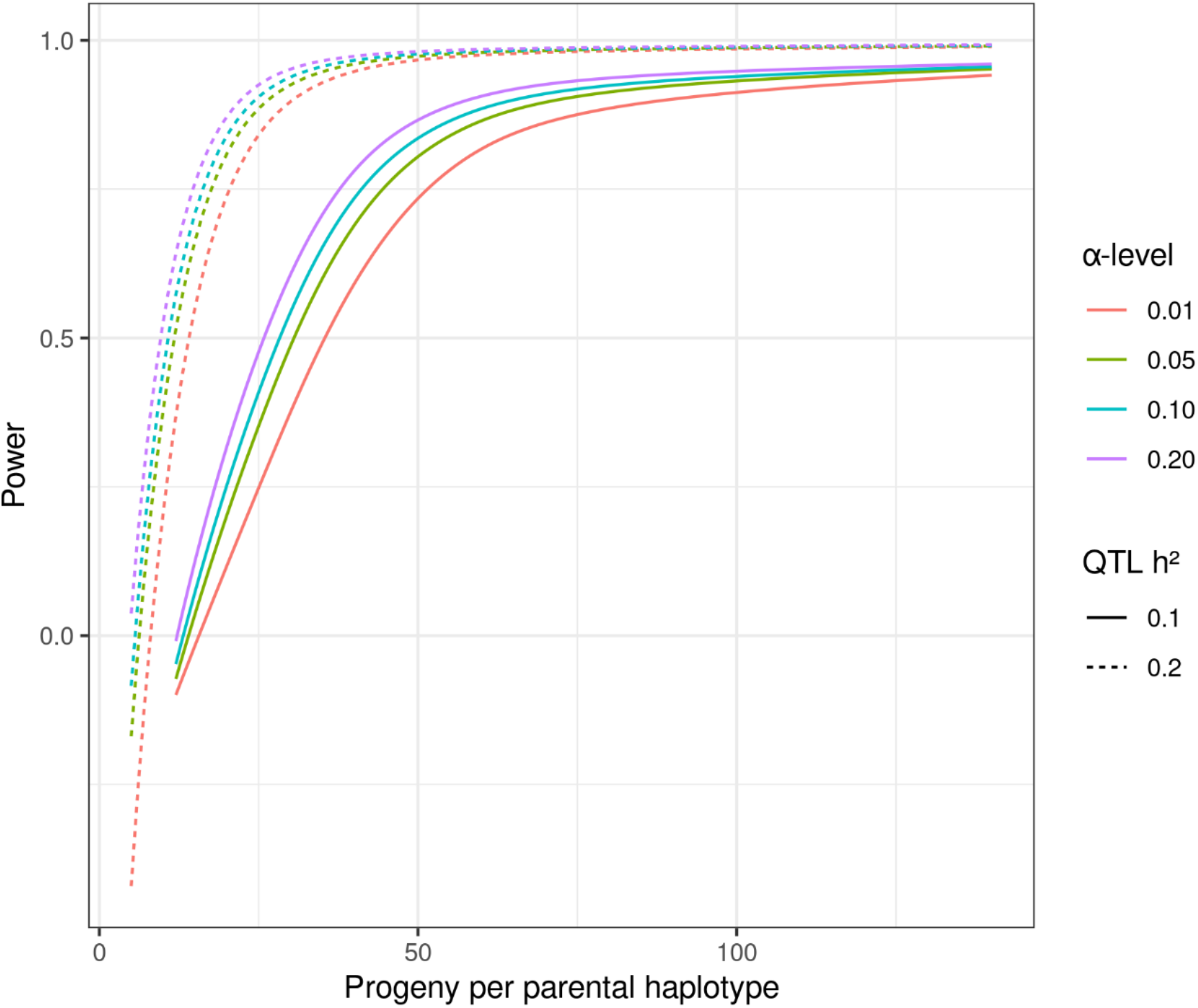
Statistical power to detect QTL for different significance levels α, as a function of the number of progeny per parental haplotype, number of parents, ploidy, and QTL heritability (h^2^). Each point is the average of 1,000 simulations with an additive model. The dashed line is a monotone increasing, concave spline for h^2^ = 0.1, and the solid line is the spline for h^2^ = 0.2.

**Figure S6.**
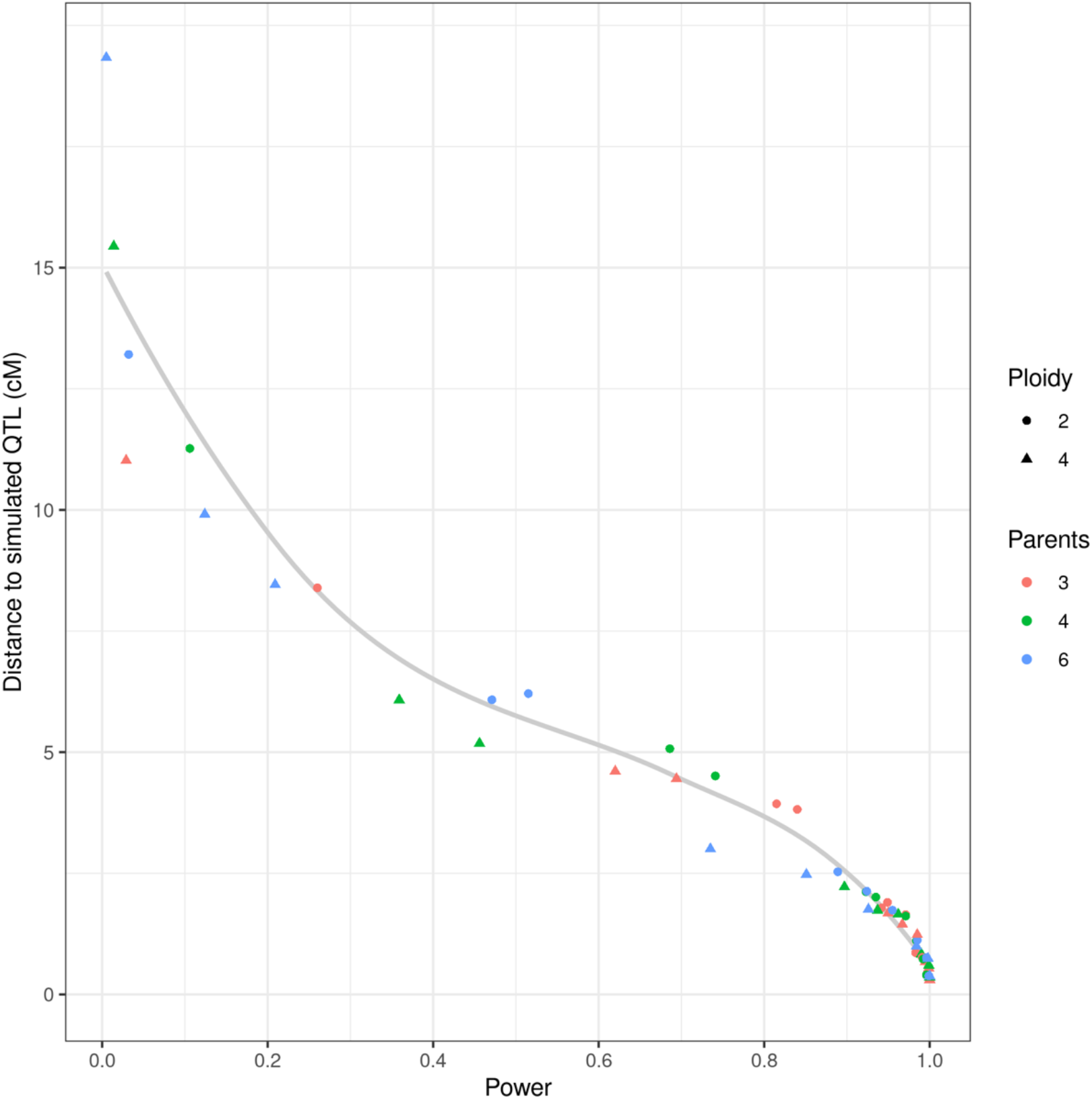
Average distance between the position of the most significant marker and simulated QTL position, as function of power, ploidy and number of parents. Each point represents an average over the multiple population sizes and heritability values shown in Figure S4.

**Figure S7.**
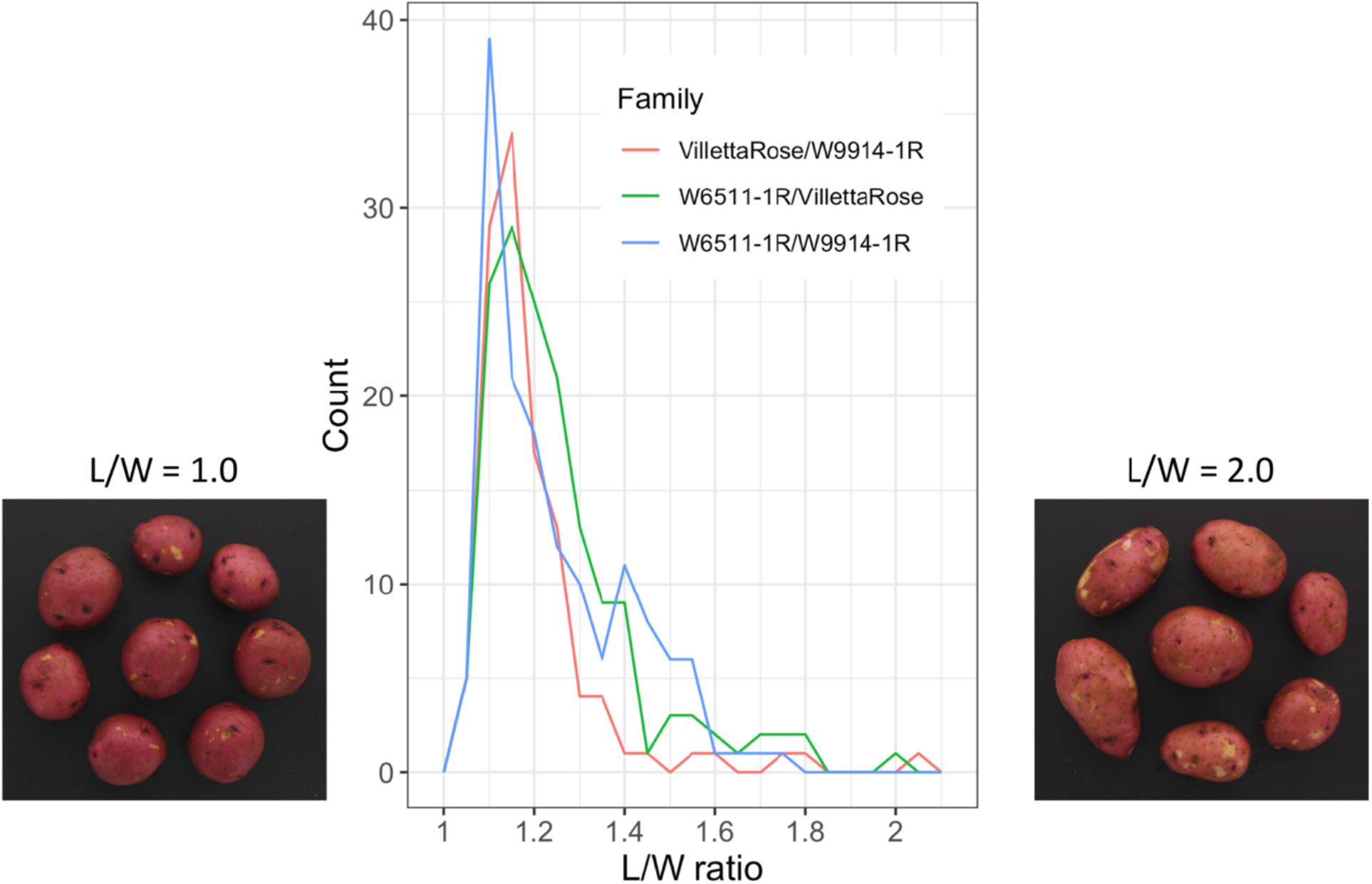
Distribution of the length to width (L/W) ratio for tubers of three full-sib families of autotetraploid potato. The images illustrate the extreme phenotypes of this population with L/W = 1.0 as a round tuber shape, L/W = 2.0 as an elongated tuber shape.

**Figure S8.**
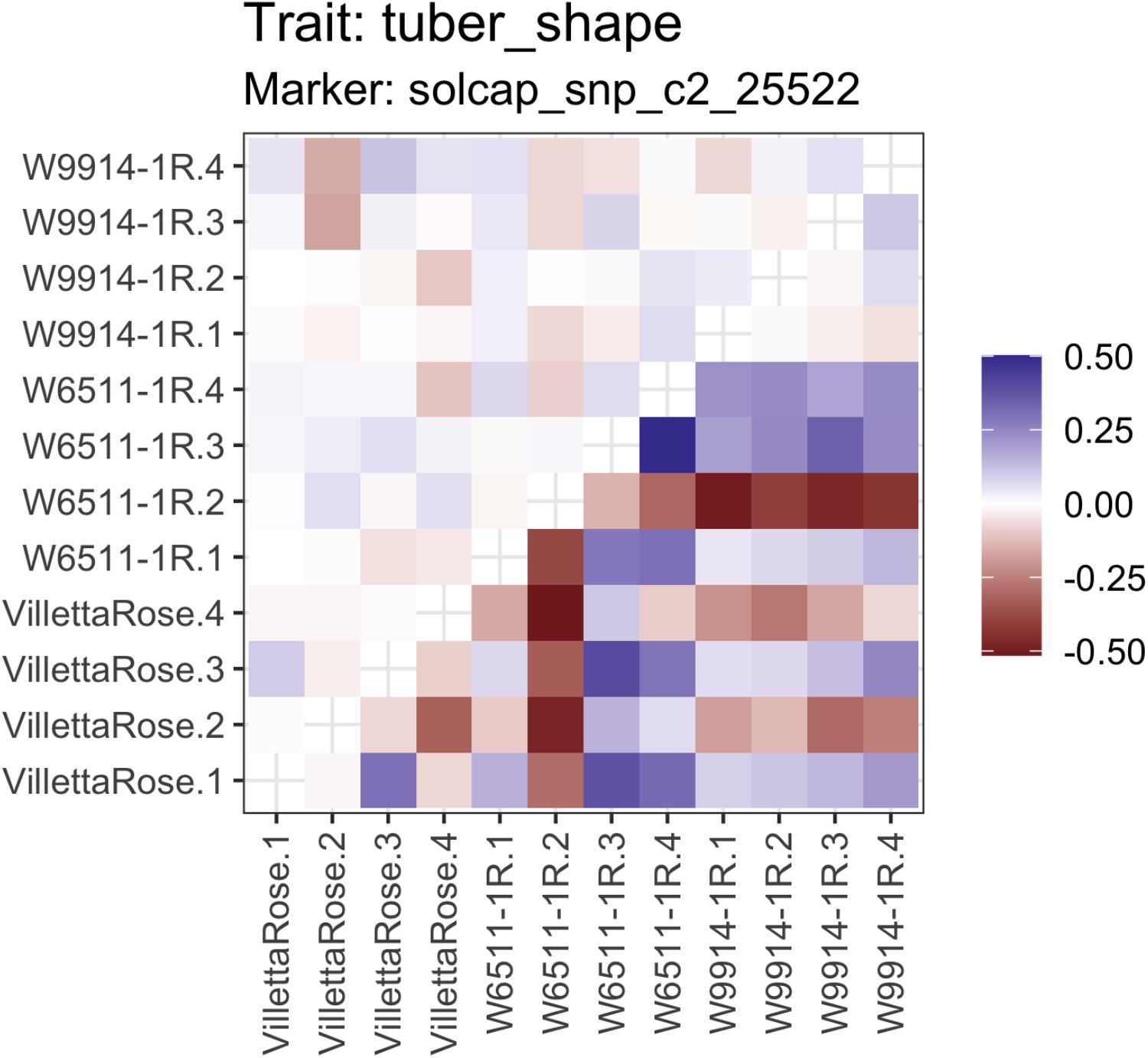
Estimated QTL effects for the 10@63 QTL with the digenic dominance model (#4 in Table 1). The digenic dominance effects are above the diagonal, and below the diagonal are the sum of the additive and digenic effects. The four haplotypes for each parent are labeled .1 to .4. The figure is part of the graphical output from diaQTL function *fitQTL*.

**Figure S9.**
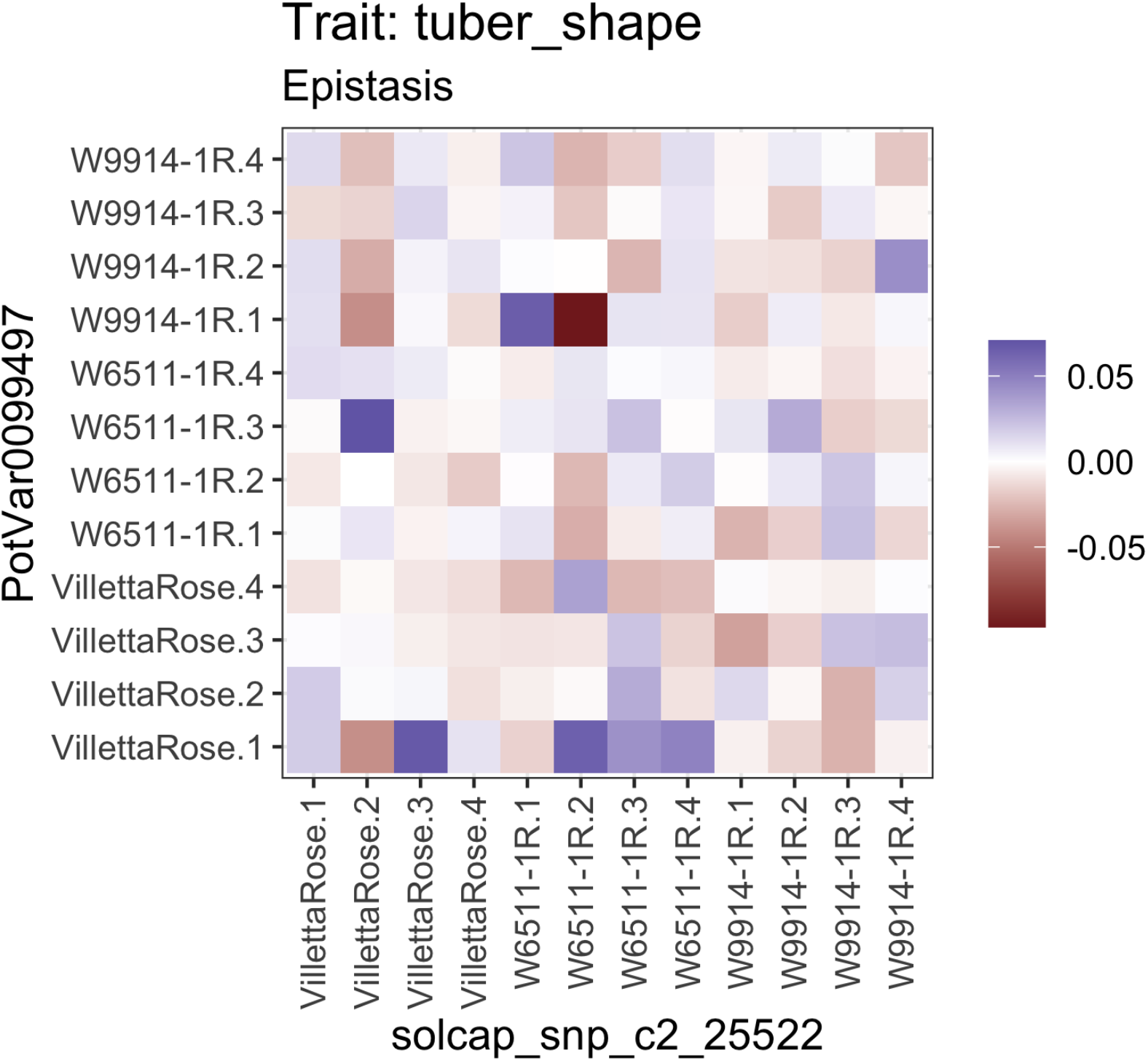
Estimated additive x additive epistatic effects between the 10@63 and 1@133 QTL (model #3 in Table 1). The four haplotypes for each parent are labeled .1 to .4. The figure is part of the graphical output from diaQTL function *fitQTL*

**Figure S10.**
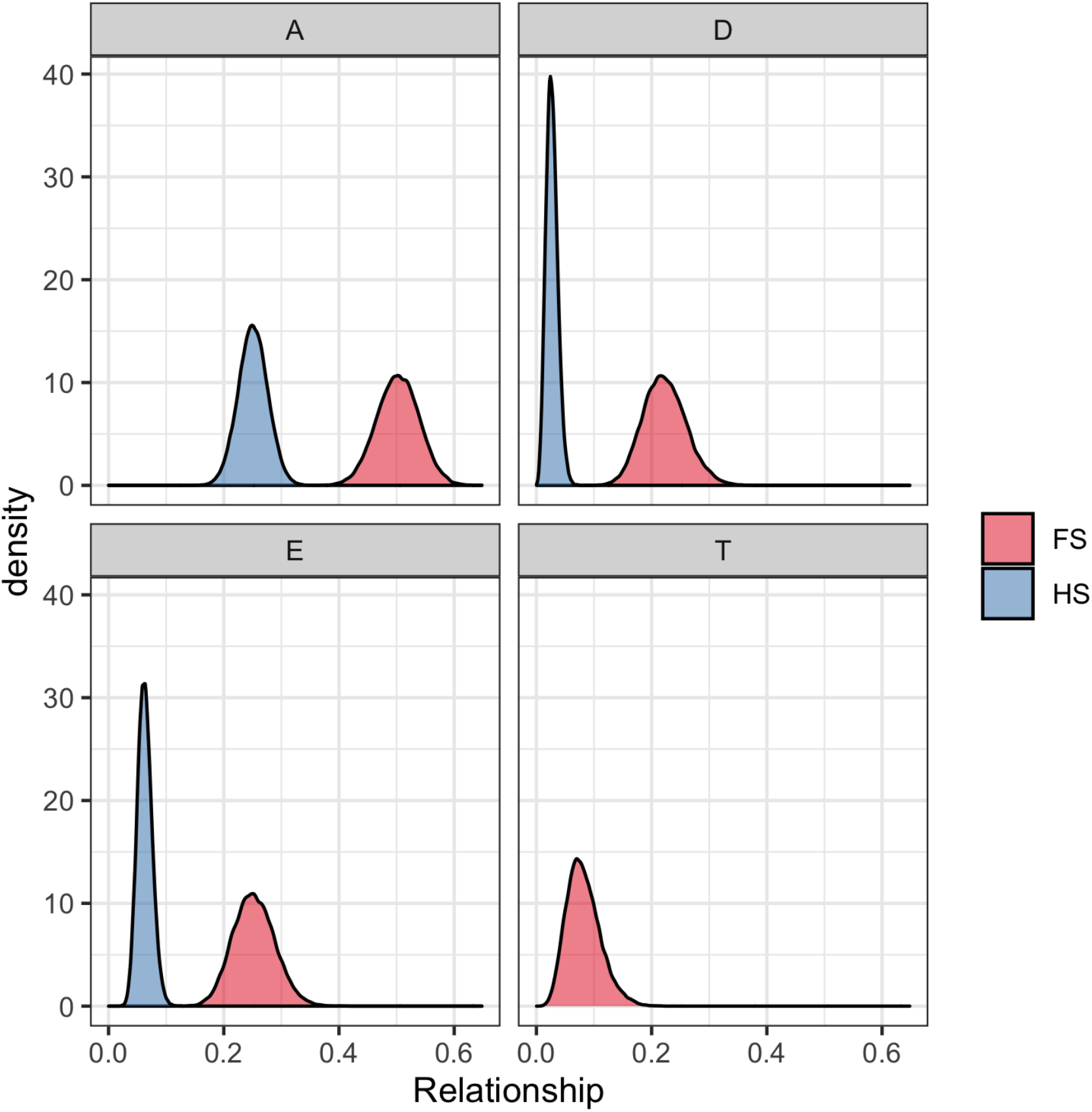
The distribution of realized relationships between full-sibs (FS) and half-sibs (HS) in the potato diallel population, computed with diaQTL function *IBDmat*. The four panels correspond to additive (A), digenic dominant (D), trigenic dominant (T), and additive x additive epistatic (E) effects. The expected values based on pedigree are A = 1/2, D = 2/9, T = 1/12, E = 1/4 for full-sibs and A = 1/4, D = 1/36, T = 0, E = 1/16 for half-sibs.

**Figure S11.**
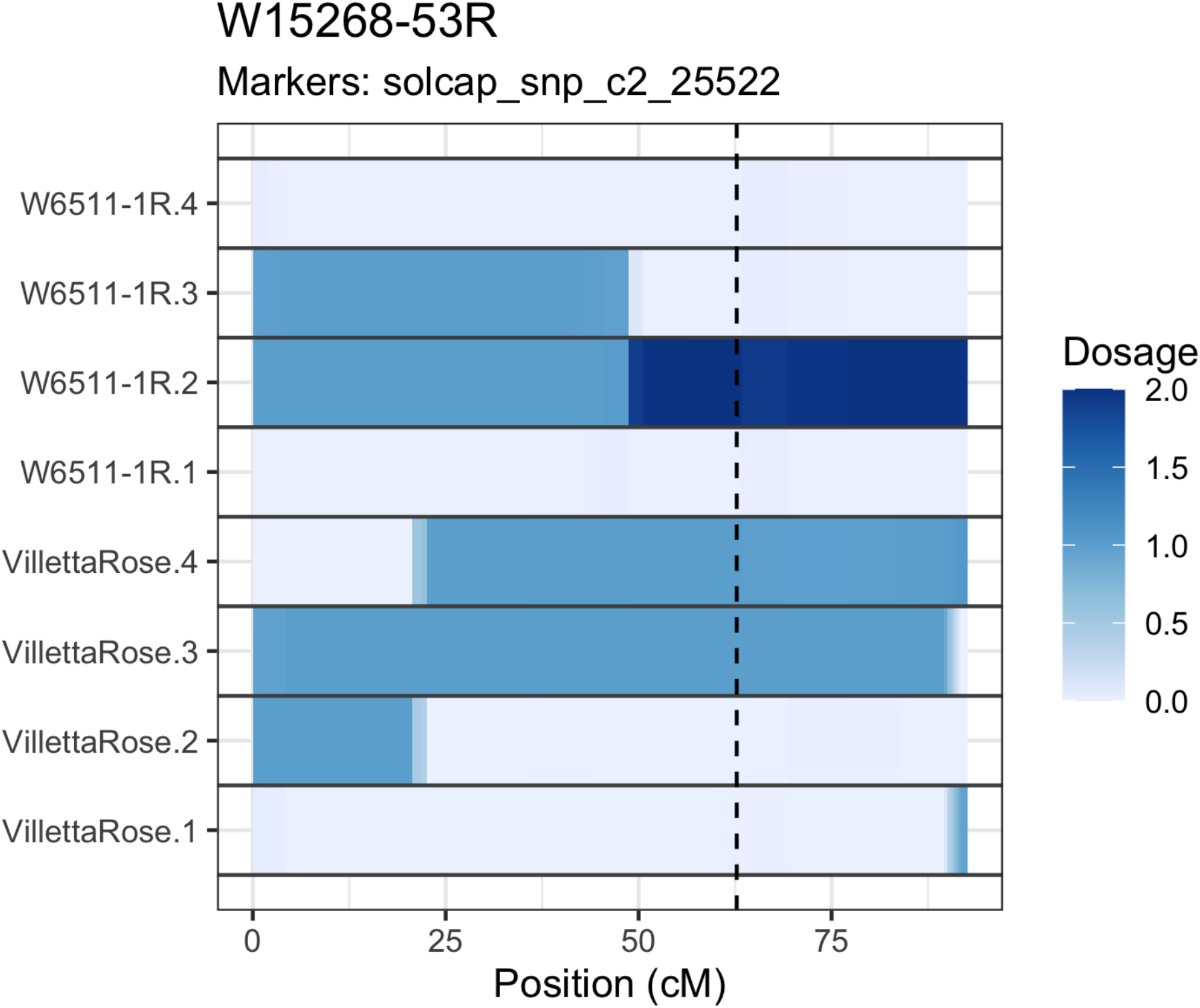
Visualizing the inheritance of parental haplotypes from VillettaRose and W6511-1R in one F1 offspring (W15268-53R), using diaQTL function *haplo_plot*. This clone was selected as one of three offspring containing two copies of the W6511-1R.2 haplotype, which was estimated to have the largest negative effect (Fig. 5) and is therefore desirable for breeding potato varieties with round tubers. The dashed line marks the position of the 10@63 QTL for tuber shape. The presence of two copies of a parental haplotype is possible in tetraploid offspring due to double reduction, a phenomenon in which the diploid gamete contains portions of both sister chromatids from one homologous chromosome.

